# Structures of an intact yeast V-ATPase alone and in complex with bacterial effector VopQ

**DOI:** 10.1101/2020.07.16.207225

**Authors:** Wei Peng, Jessie Fernandez, Amanda K. Casey, Lisa N. Kinch, Diana R. Tomchick, Kim Orth

## Abstract

Vacuolar-type ATPase (V-ATPase) is a rotary protein pump involved in proton translocation across various cellular membranes using the energy of ATP hydrolysis. Despite previous studies on bacterial and eukaryotic V-ATPases, information on the intact structure of a eukaryotic V-ATPase is missing. Here we report cryo-EM structures of the intact yeast V-ATPase and this complex bound to a bacterial effector. We reveal the interaction of the elusive regulatory subunit H with its neighboring subunits. Insight for the catalysis mechanism is gained by determining conformations of the catalytic subunits either empty or bound with nucleotides.

## Introduction

V-ATPase is a giant protein complex with the primary function of pumping protons into organelles to create a local acidic environment and membrane potential for biological processes such as protein degradation in lysosome^1^. As an important protein machine present in various membranes, V-ATPase is also involved in other cellular functions exemplified by mTOR signaling, Notch signaling, neurotransmitter release, and bone remodeling^2–5^. Consequently, dysfunction of V-ATPase is associated with related diseases.

V-ATPase is composed of 16 known unique subunits in yeast and mammals with 31 and 32 polypeptides, resulting in a molecular mass of around 1 megadalton^6–8^. The proton translocation is built on the coupling between ATP hydrolysis within the cytosolic V_1_ subcomplex and proton transport within the membrane-inserted V_o_ subcomplex^9,10^. The V_1_ subcomplex contains subunits of A_3_, B_3_, C, D, E_3_, F, G_3_, and H and the V_o_ subcomplex includes subunits of a, c_8_, c′, c″, d, e, f, and Voa1 in yeast or a, c_9_, c″, d, e, f, Ac45, and PRR in mammals^6–8^. As a member of the rotary ATPases, V-ATPase shares similarities in the architecture and working mechanism with F-ATPase (F-type) and A-ATPase (archaeal type)^11–13^. However, V-ATPase still differs in many ways from the well-studied F-ATPase or A-ATPase that usually catalyze ATP synthesis driven by protonmotive force. Major differences in V-ATPase include three copies of peripheral stalk subunits (E_3_G_3_) instead of one (or two in A-ATPase) and additional subunits of C and H, larger c-ring subunit scaffold (two helix hairpins containing four helices), regulated disassembly in response to environmental factors, and the catalysis of ATP hydrolysis instead of ATP synthesis^10,11^.

Recent progress on bacterial V/A-ATPases and eukaryotic V-ATPases has begun to shed light on the structures and mechanisms of V-ATPase^7,8,14–17^. Although structures of a rat V-ATPase were reported recently^8^, the rat V-ATPase is not an intact holoenzyme as it lacks the regulatory subunit H. Herein, we describe cryo-EM structures of the first intact eukaryotic holoenzyme V-ATPase complex (V_1_V_o_) and this holoenzyme in complex with a bacterial effector (V_1_V_o_-VopQ). Using high resolution structures containing bound nucleotides (AMP-PNP and ADP), we illustrate the mechanism of enzyme catalysis in a V-ATPase holoenzyme complex.

## Results

### Structure determination of the intact yeast V-ATPase holoenzyme

Yeast V-ATPase contains two isoforms of subunit a (Vph1p and Stv1p) and only one isoform of all the other subunits^6,18,19^. Therefore, to obtain a homogeneous yeast V-ATPase complex with the single subunit a isoform (Vph1p), we deleted Stv1p. This complex was supplemented with an ATP analog, AMP-PNP, in order to capture different conformations of V-ATPase and stabilize the complex by using this hydrolysis-deficient substrate. Cryo-EM data analysis indicates the presence of state 1 and state 2 complexes, but not state 3, with the majority of particles in state 1 (**Extended Data Fig. 1**). We propose that state 3 likely disassembles during the purification steps, consistent with the V-ATPase disassembly model that state 3 is where disassembly starts^15,20,21^. The V_1_V_o_ complex obtained was intact with subunit H density maps resolved.

State 1 of the V_1_V_o_ complex was determined at a reported overall resolution of 3.14 Å, with focused refinements further improving local resolutions (**Extended Data Fig. 1,3,4**). State 2 was resolved at reported overall resolution of 3.53 Å, however, the local density maps were worse due to the problem of preferred orientation (**Extended Data Fig. 3d**). The yeast V_1_V_o_ protein sample was supplemented with AMP-PNP causing closure of the catalytic subunit pair of A_(1)_-B_(1)_ (**Fig. 1**). A gray line in the model is drawn through the A-B subunit pairs to draw attention their open and closed states. Previously, this same pair was shown to be in a loose conformation in nucleotide free environment^15^. The catalytic subunit pair of A_(2)_-B_(2)_ are in an open conformation. The third pair, A_(3)_-B_(3)_ is in a closed state and occupied by an AMP-PNP molecule that replaces ADP as was modelled in the rat V-ATPase^8^.

**Fig. 1.**
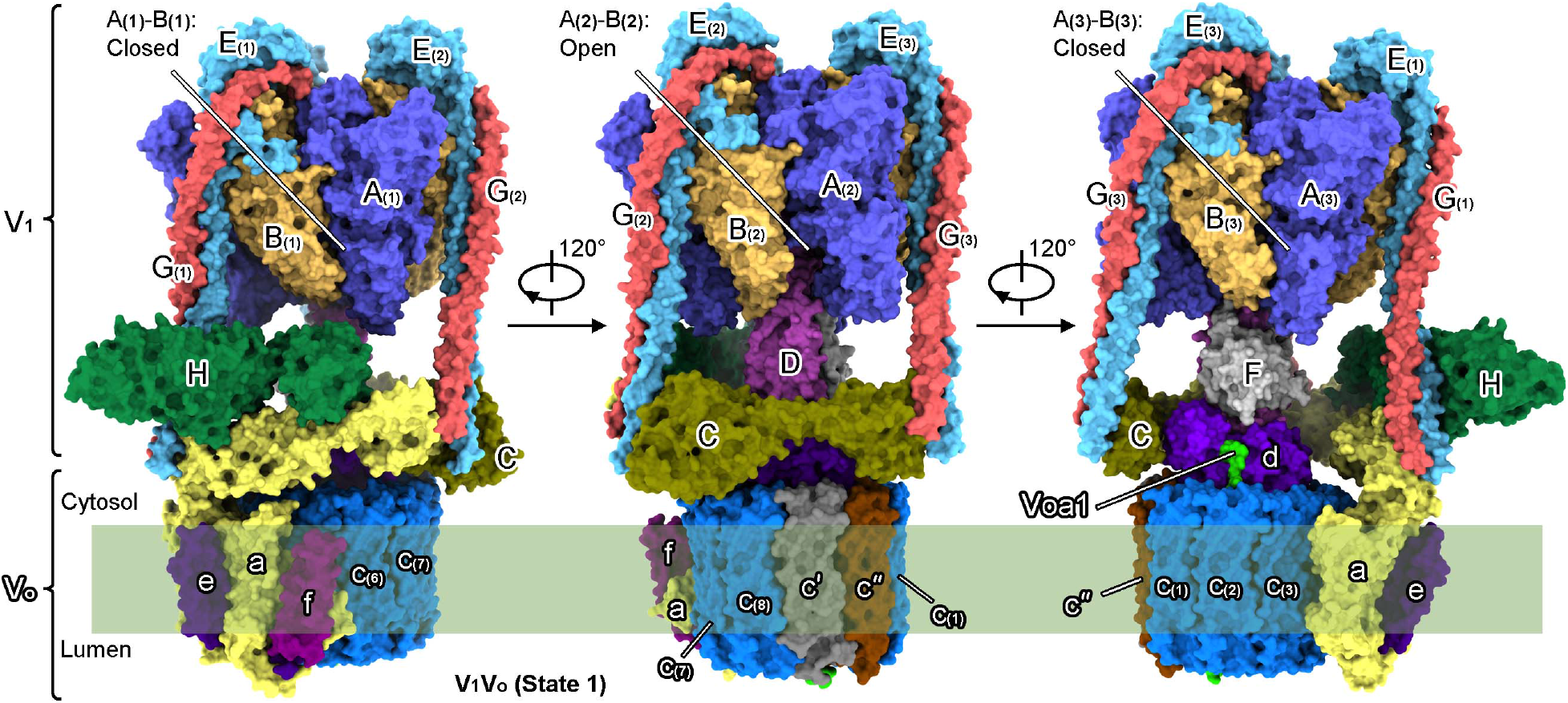
Structure of the yeast V_1_V_o_ holoenzyme in state 1. Surface presentation of the yeast V_1_V_o_ holoenzyme in state 1 shown with each individual subunit colored and indicated. V_1_ subunits (in solid uppercase) include three copies of subunit A, B, E, and G, and one copy of subunit C, D, F, and H. V_o_ subunits (in hollow lowercase) include one copy of subunit a, c′, c″, d, e, f, and Voa1, and eight copies of subunit c. All subunits are colored using the same color scheme in all figures unless otherwise noted. The catalytic subunit pairs of A_(1)_-B_(1)_ and A_(3)_-B_(3)_ are in closed state while A_(2)_-B_(2)_ is open.

### V_1_V_o_ holoenzyme in complex with a bacterial effector

A *Vibrio parahaemolyticus* type 3 secretion system effector, VopQ, was previously shown to bind dissociated V_o_ subcomplex and predicted to bind the holoenzyme of V_1_V_o_ in state 2 (ref. ^17^). To gain better insights into V_1_V_o_ and test if VopQ could indeed interact with the holoenzyme, we prepared the V_1_V_o_ sample with VopQ present and analyzed the cryo-EM data (**Extended Data Fig. 2,3**). Both AMP-PNP and ADP were supplemented in the V_1_V_o_-VopQ sample to obtain ADP bound V-ATPase. As previously predicted, V_1_V_o_-VopQ complex was found to be exclusively in state 2 and the angular distribution was restored to 3 major preferred orientations, similar to the V_1_V_o_ complex in state 1 (**Extended Data Fig. 3d**). This observation suggests that VopQ interacts with V_1_V_o_ in state 2 and locks the enzyme in this state. The final 3D reconstruction improved the cryo-EM density map quality and allowed characterization of V_1_V_o_ in state 2 (**Extended Data Fig. 4**).

A detailed examination and comparison of the 3D class cryo-EM density maps indicated that VopQ was bound with V_1_V_o_ in state 2 as predicted (**Extended Data Fig. 5a**). However, the density maps for VopQ were not well resolved, indicating that VopQ may partially clashed with subunits in V_1_V_o_ and the relatively dynamic V_o_ regions may also contribute to this clash. A VopQ molecule corresponding to VopQ-2B in the reported V_o_-VopQ structures^17^ was docked into the cryo-EM density map (**Fig. 2a**). To further support the hypothesis that VopQ interacts with V_1_V_o_ in state 2, we performed an ATPase activity inhibition assay (**Extended Data Fig. 5b**). VopQ showed no inhibition on ATP hydrolysis when ATP was added before V_1_V_o_. But when VopQ was pre-incubated with V_1_V_o_ before ATP was added, a partial inhibition of ATP hydrolysis was observed. This result suggests that the interaction between VopQ and V_1_V_o_ requires V_1_V_o_ to be relatively stable in state 2 so VopQ can bind while changing from a soluble state to a membrane imbedded and V_1_V_o_-bound state. Under both conditions, the V-ATPase inhibitor folimycin^22^ could completely block ATP hydrolysis, implying a rapid binding of this inhibitor. The complete inhibition by folimycin indicates a tight coupling between V_1_ ATP hydrolysis and V_o_ proton transport as it works at the transmembrane region of V_o_ subcomplex^23^.

**Fig. 2.**
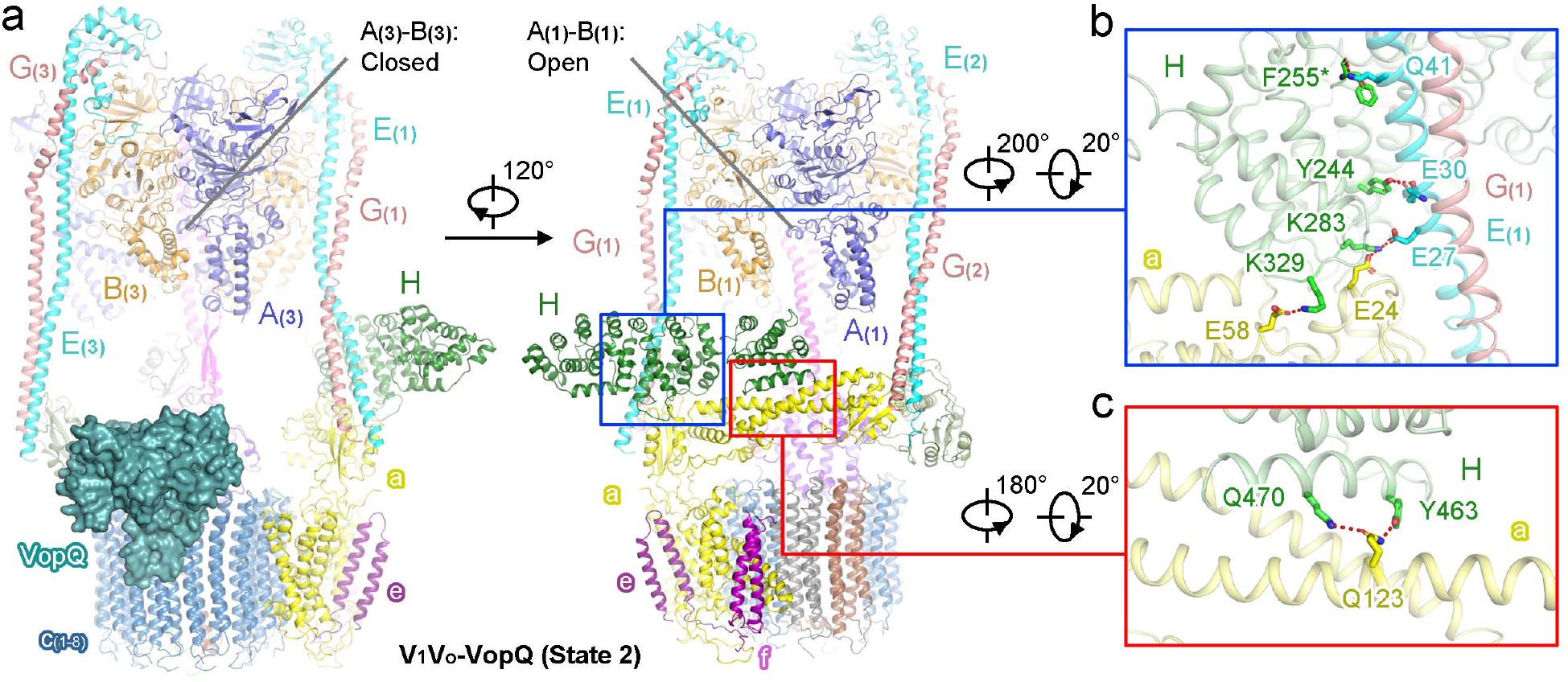
Structure of the yeast V_1_V_o_-VopQ complex in state 2. **a**, The yeast V_1_V_o_-VopQ complex in state 2. VopQ is shown as surface. Subunit H is in close contact with V_1_ subunit E_(1)_, G_(1)_, and V_o_ subunit a. **b**,**c**, Interaction between subunit H and subunit G or subunit a. Residues involved in hydrogen bond or salt bridge interaction are shown as sticks. The main chain carbonyl oxygen atom of F255 in subunit H, indicated by “*”, forms a hydrogen bond with Q41 in subunit E_(1)_. Both the side chain and main chain oxygen atoms of E30 in subunit E_(1)_ are involved in the interaction with Y244 in subunit H. All other interaction pairs are mediated by sides chains as indicated.

Disassembled V_1_ will continuously drive ATP hydrolysis without subunit H which plays an inhibitory role^24^. This activity will not be inhibited by V_o_ inhibitors (folimycin or bafilomycin)^23^ as was observed for the rat V-ATPase that lacks subunit H^8^. Therefore, our data also suggests that subunit H is present in possible disassembled V_1_ subcomplex and in the holocomplex, supported by the fact that subunit H is essential for the holoenzyme activity^25^.

In the V_1_V_o_-VopQ (state 2) structure, conformations of catalytic subunit pair A_(3)_-B_(3)_ (closed), A_(1)_-B_(1)_ (open), and A_(2)_-B_(2)_ (closed) correspond to those of A_(1)_-B_(1)_, A_(2)_-B_(2)_, and A_(3)_-B_(3)_ in state 1 due to enzyme reaction and rotation of rotor subunits (**Fig. 2a**). The catalytic site of the A_(2)_-B_(2)_ pair is bound with an ADP molecule rather than an AMP-PNP molecule. The overall conformations of catalytic subunit pairs are similar to those in the previously published disassembled V_1_ (not shown), which is in an inhibited state similar to state 2 (ref. ^26^).

### Interaction of subunit H with the other V-ATPase subunits

Subunit H is a regulator that inhibits ATP hydrolysis by disassembled V_1_ and interestingly, subunit H is necessary for the activity of the holoenzyme^24,25,27^. In the yeast holoenzyme obtained above, although subunit H was present, the intra-complex dynamics between V_1_ and V_o_ subcomplexes hindered analysis of the interaction between subunit H and neighboring subunits. The V_1_V_o_-VopQ complex displayed better map quality for subunit H and allowed definition of this interaction, possibly stabilized by the binding of the protein inhibitor VopQ (**Extended Data Fig. 4**).

The N-terminal domain of subunit H is in close contact with V_1_ subunit E_(1)_, subunit G_(1)_, and V_o_ subunit a, while the C-terminal domain is contacting V_o_ subunit a but not the V_1_ subcomplex (**Fig. 2a**). Density-supported interaction pairs include (* indicates main chain atom): Y244-E30_E_(1)_, Y244-E30*_E_(1)_, F255*-Q41_E_(1)_, K283-E27_E_(1)_, K283-E24_a, and K329-E58_a (N-terminal, **Fig. 2b**); Y463-Q123_a and Q470-Q123_a (C-terminal, **Fig. 2c**). These interactions are in contrast to the yeast V-ATPase V_1_ subcomplex where the C-terminal domain is folded up toward the V_1_ subunits B_(1)_ and D^26^. Our structures support the previous model where C-terminal domain of subunit H is flexible and is able to swing out when the V_1_ and V_o_ domains join to form a holoenzyme^26^. Numerous cancer-related mutations can be mapped to human V-ATPase subunits when searched in BioMuta^28^. A few of these mutations are located near the interfaces between subunit H and subunit E_(1)_, G_(1)_, and a. In the yeast V-ATPase, some of the corresponding residues are: R229, E284, S324, R328, D332, T469, Q470 (subunit H); F23, I24, E27 (subunit E); T290, T291, T294 (subunit a). These mutations possibly cause dysfunction of V-ATPase by disrupting the regulation activity of subunit H.

### Conformations of the catalytic subunit pairs

As mentioned above, A_(1)_-B_(1)_, A_(2)_-B_(2)_, and A_(3)_-B_(3)_ in state 1 are in closed, open, and closed conformations. The A_(1)_-B_(1)_ in state 1 is captured in a closed conformation due to the binding of non-hydrolysable AMP-PNP (**Fig. 3a**). The natural product of ATP hydrolyses, ADP, is modelled in the rat V-ATPase^8^ A_(3)_-B_(3)_ pair and is replaced in this structure by AMP-PNP, fitting well in the density map (**Extended Data Fig. 4c**). In the V_1_V_o_-VopQ complex in state 2, similar conformational arrangement is observed for A_(1)_-B_(1)_, A_(2)_-B_(2)_, and A_(3)_-B_(3)_ with pairs in the open, closed, and closed conformations, respectively. However, the A_(2)_-B_(2)_ catalytic site for the V_1_V_o_-VopQ complex is occupied by an ADP molecule instead of AMP-PNP (**Fig. 3b; Extended Data Fig. 4c**). As both ADP and AMP-PNP were supplemented at equal concentration to the final sample, this might support the proposal that ADP has a higher affinity at this site than AMP-PNP. Conversely, AMP-PNP might have a higher affinity at the A_(3)_-B_(3)_ catalytic site. The two catalytic sites in the V_1_V_o_ in state 2 without VopQ, where ADP is not provided, are both occupied by AMP-PNP supported by the density maps (**Extended Data Fig. 4c**).

**Fig. 3.**
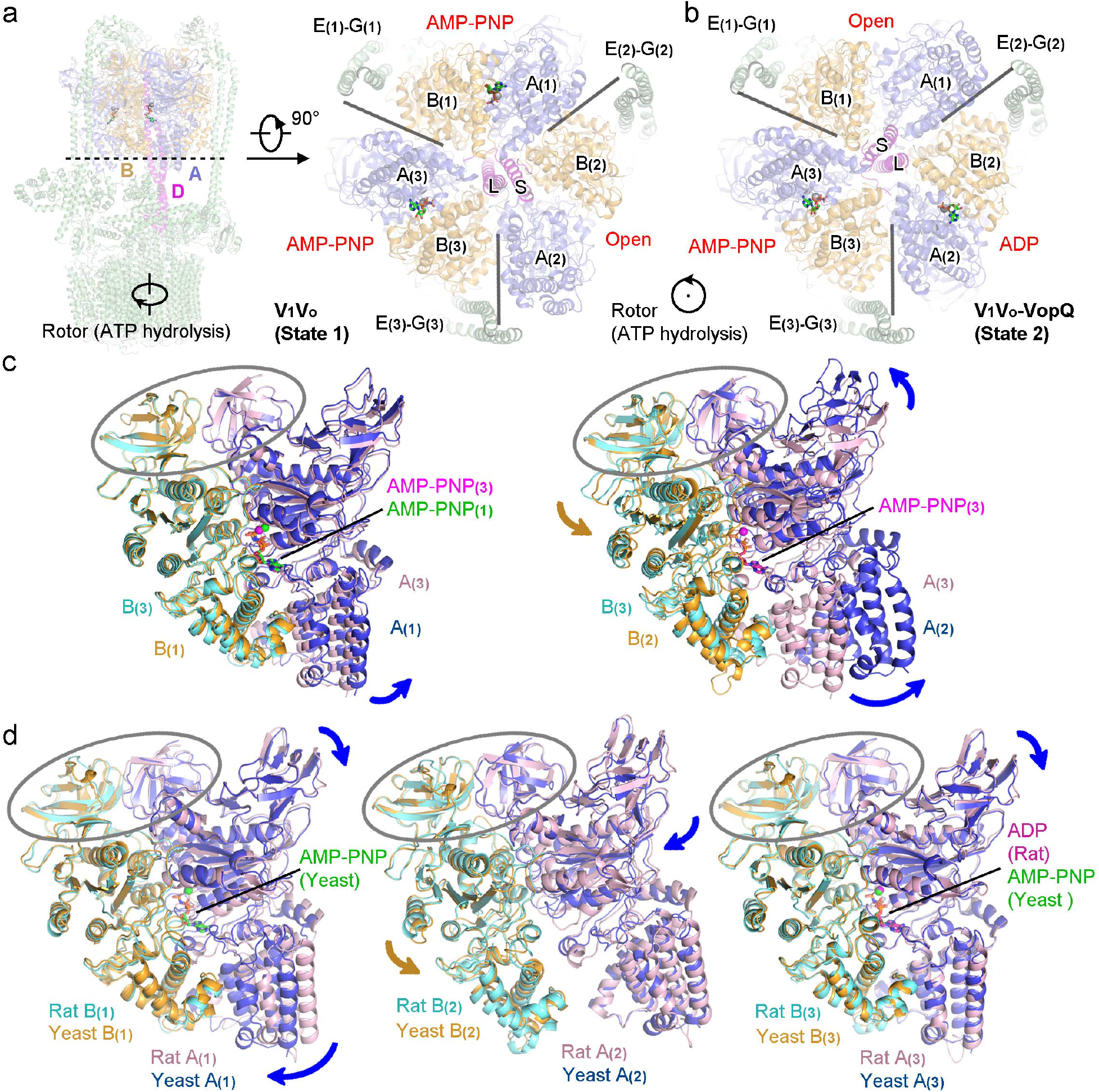
Conformations of yeast V_1_V_o_ catalytic subunit pairs. **a,b**, The yeast V_1_V_o_ complex in state 1 and the yeast V_1_V_o_-VopQ complex in state 2. The three catalytic subunit pairs are indicated and separated by radial lines. Subunit D and catalytic subunits are colored and indicated with all the other subunits in green. Nucleotide molecules are shown as sticks with magnesium ions as spheres. “S” or “L”, short N-terminal helix or long C-terminal helix of subunit D. **c**, Comparison of subunit A-B pairs of the yeast V_1_V_o_ complex in state 1. **d**, Comparison of subunit A-B pairs in the yeast V_1_V_o_ complex (state 1) and in the rat complex (state 1). Catalytic pairs are aligned against the N-terminal β-barrel domains indicated by ovals. Relative motions are indicated by colored arrows.

V-ATPase is a member of rotary ATPases that also include F-ATPase and A-ATPase that can both synthesize and hydrolyze ATP^11–13^. Rotary ATPase are dynamic protein machines that have been captured in many states and conformations, either for the subcomplexes or the holoenzymes. Rotary ATPases share similarities in architectures and working mechanisms (**Extended Data Fig. 6a**). When visualizing the catalytic subunits from the membrane side in representative rotary ATPase structures and placing the short N-terminal helix of subunit γ in F-ATPase or subunit D in V-ATPase on the left and the long C-terminal helix on the right (**Extended Data Fig. 6b-6e**), the top catalytic pairs are in open and therefore empty conformations. This regular pattern also applies to our structures (**Fig. 3a,3b**). By comparing our structures with and a bacterial intact V/A-ATPase and other eukaryotic V-ATPase structures (V_1_ or V_1_V_o_) based on this pattern, it is obvious that yeast V-ATPase structures are in different states with two of the three catalytic A-B pairs occupied by nucleotides^8,15,16^ (**Fig. 3a,3b; Extended Data Fig. 6d,6e**).

V-ATPase catalytic subunit (A or B) can be roughly divided into three domains, an N-terminal β-barrel domain, a central nucleotide-binding domain (for subunit A, an additional non-homologous domain), and a C-terminal domain^29^ (**Extended Data Fig. 7a**). The conformations of each catalytic subunit in V_1_V_o_ (state 1) is examined and compared (**Extended Data Fig. 7b**). Subunit A and B in A_(1)_-B_(1)_ pair are almost identical to those in A_(3)_-B_(3)_ pair, resulting in small rigid movement of the A_(1)_-B_(1)_ pair compared to A_(3)_-B_(3)_ when aligned against the N-terminal β-barrel domains (**Fig. 3c**). Subunit A or B in A_(2)_-B_(2)_ pair undergoes obvious intra-subunit conformations changes, contributing to their open conformation (**Fig. 3c; Extended Data Fig. 7b**). In comparison with the rat V-ATPase in state 1, the yeast A_(1)_-B_(1)_ pair shows an obvious more closed conformation while the other two pairs are more or less similar (**Fig. 3c; Extended Data Fig. 7c**). The A_(3)_-B_(3)_, A_(1)_-B_(1)_, and A_(2)_-B_(2)_ pairs in V_1_V_o_-VopQ of state 2 are in similar intra-subunit and subunit-pair conformations with A_(1)_-B_(1)_, A_(2)_-B_(2)_, and A_(3)_-B_(3)_ pairs in V_1_V_o_ of state 1 (**Extended Data Fig. 8**). These observed conformations represent pre-catalysis states of the catalytic subunits in the holoenzyme, captured using non-hydrolysable AMP-PNP.

## Discussion

In the yeast V-ATPase structures described above, extended residues of subunit B (N terminus) and subunits E and G (C terminus) were modelled compared to the previous low-resolution structures^15^. These residues indicate additional interaction interfaces between subunit B, E, and G in the yeast V-ATPase as described for the rat V-ATPase^8^. The bacterial effector VopQ is capable of binding the disassembled V_o_ subcomplex^17^. As *in silico* predicted, VopQ helps to resolve the V-ATPase structure in state 2 as a state-specific inhibitor, making it a useful tool for future V-ATPase studies.

Eukaryotic V-ATPase is a member of rotary ATPases that differs from F-ATPase and A-ATPase. With the high-resolution intact yeast V-ATPase structures, we are able to visualize the proton pump with subunit H assembled which is an important regulator of eukaryotic V-ATPase. Studies suggest that subunit H prevents energy waste of ATP hydrolysis in disassembled V_1_ subcomplex^24,26^. In the V-ATPase holoenzyme, it is necessary for activity but not assembly^25,27^. Since subunit H only contacts V_1_ subunits E_(1)_ and G_(1)_ and V_o_ subunit a in the holoenzyme, it is likely that subunit H holds the framework of stator subunits and allows ATP hydrolysis by the catalytic subunits. Though the reported rat V-ATPase structures without subunit H^8^ show differences in the stator subunits compared to our yeast structures (not shown), it is not known if these differences result from sample variety. Therefore, systematic studies of a eukaryotic V-ATPase from a single organism is necessary for revealing possibly subtle differences between different states or conformations, which will contribute to an understanding of subunit H function. This calls for structures of yeast V-ATPase without subunit H or structures of rat V-ATPase with subunit H. The yeast V-ATPase represents a better option as extensive studies have been performed on it and it is easier to manipulate the much less genes of subunit isoforms in yeast for biophysical and biochemical tests.

Catalysis mechanisms for rotary ATPases have been proposed mostly based on F_1_ and V_1_ subcomplexes^13,14,29^. However, these subcomplexes only contain the core catalytic subunits and rotor subunits, with some even lacking the rotor subunits^29^. The incompleteness of these subcomplexes will cause structure feature changes compared to the holoenzymes as observed^14,29^. These changes are likely artificial. Even the disassembled intact yeast V_1_ subcomplex, which is physiologically related, is locked in an auto-inhibited conformation that differs from the holoenzyme^26^. Therefore, direct observation of various conformations of catalytic subunit pairs in a V-ATPase holoenzyme was needed to test the rotary catalysis mechanism. By determining the conformations of catalytic subunit pairs in the yeast V-ATPase holoenzyme, we revealed two similar nucleotide-bound conformations and one open conformation. These observations in a V-ATPase holoenzyme support the previous model^13,14,30^ of a rotary catalysis mechanism wherein: (1) two catalytic subunit pairs bind ATP molecules with one being empty; (2) the ATP bound pair after the empty in the rotation direction catalyzes ATP hydrolysis; (3) ADP and phosphate get released and the empty pair is occupied by an ATP molecule; (4) the catalytic subunit pairs repeat the above steps. Therefore, our studies provide a mechanistic insight into a fully assembled and functional V-ATPase and are helpful for future studies to address other rotary ATPases.

## Acknowledgments

We thank X. Li and the Orth lab members for discussions and editing. We thank the Friedman laboratory for assistance with yeast cell disruption. We thank the Structural Biology Laboratory (SBL) for assistance with cryo-EM grid preparation and screening. Cryo-EM data were collected at the University of Texas Southwestern Medical Center Cryo-Electron Microscopy Facility that is funded in part by the CPRIT Core Facility Support Award RP170644. We thank SBL for providing computational resources for cryo-EM data processing. This work was funded by the Welch Foundation grant I-1561 (K.O.), Once Upon a Time…Foundation (K.O.), and National Institutes of Health Grant R01 GM115188 (K.O.). K.O. is a W.W. Caruth, Jr. Biomedical Scholar with an Earl A. Forsythe Chair in Biomedical Science. D.R.T. is an Effie Marie Scholar.

## Author contributions

W.P. and K.O. conceived the project and designed the experiments. W.P., J.F., A.K.C., and L.N.K. conducted the experiments. W.P. and K.O. prepared the manuscript with input from all authors. K.O. supervised the project.

## Methods

### Purification of the Yeast V-ATPase Complex

The *S. cerevisiae* TAP-tagged *VMA1* strain (YSC1178-202230289, Dharmacon, Inc.) was purchased and the TAP tag was replaced by a 3×FLAG tag after *STV1* was knocked out^31^. Yeast cells (8~9 liters in flasks) were cultured to logarithmic phase which helped maintain an intact holoenzyme because of abundant glucose^32^. Collected cells were resuspended in lysis buffer (25 mM HEPES pH 7.4, 150 mM NaCl, 5 mM EDTA, 1 mM PMSF, protease inhibitor cocktail, 2% glucose, 8% sucrose, 2% sorbitol) and frozen in liquid nitrogen as dropped beads (diameter less than 3 mm). The frozen cells were disrupted using a 6870D Freezer/Mill^®^ Dual Chamber Cryogenic Grinder (SPEX™ SamplePrep). The powder thawed and the cell debris was removed by low-speed centrifugation (1,250 g, 10 min). V-ATPase containing membrane was collected by high-speed centrifugation (27,000 g, 1h). The membrane was solubilized with detergent mixture (2% n-Dodecyl-β-D-Maltoside (DDM), 0.5% 3-((3-Cholamidopropyl) Dimethylammonio)-1-Propanesulfonate (CHAPS), 0.05% Cholesteryl Hemisuccinate Tris Salt (CHS)). Insoluble materials were removed by high-speed centrifugation. The supernatant was loaded into gravity column containing anti-FLAG M2 affinity gel (GE Healthcare). The resin was washed with wash buffer (25 mM HEPES pH 7.4, 150 mM NaCl, 1 mM PMSF, protease inhibitor cocktail, 0.05% glyco-diosgenin (GDN)). Bound protein was eluted with elution buffer (25 mM HEPES pH 7.4, 150 mM NaCl, 1 mM PMSF, protease inhibitor cocktail, 0.05% GDN, 100 μg/mL 3×FLAG peptide (Sigma)). The eluted protein sample was concentrated and applied to gel filtration (Superose 6 10/300 GL, GE Healthcare) in running buffer (25 mM HEPES pH 7.4, 150 mM NaCl, 2mM DTT, 0.05% GDN). Fractions containing the V-ATPase complexes were pooled together and concentrated to around 40 μL (less than 2 mg/mL).

For preparation of V_1_V_o_-VopQ complex, VopQ protein was obtained as previously described^17^. 0.9 mg of VopQ protein was added at the membrane solubilization step. 0.2 mg of VopQ protein was added again after the complex eluted from the anti-FLAG column. The detergent used after membrane solubilization was 0.05% digitonin instead of GDN.

For cryo-EM sample preparation, 2 mM MgCl_2_ and 0.2 mM AMP-PNP were added into the V_1_V_o_ sample. 2 mM MgCl_2_, 0.2 mM AMP-PNP, and 0.2 mM ADP were added into the V_1_V_o_-VopQ sample before final concentration.

### Malachite Green Assay

Malachite green assay was performed as described with some modifications^33^. Briefly, malachite green oxalate (13.5 mg) was dissolved in 30 mL of Milli-Q H_2_O. 10 mL of 4.2% (w/v) ammonium molybdate in 4M HCl was then added. This solution was incubated on a nutator at 4 °C for at least 45 minutes. The solution was filtered and 0.01% Tween 20 was added before use.

The malachite green assay buffer contained 25 mM HEPES (pH 7.4), 150 mM NaCl, 2 mM MgCl_2_, and 0.02% DDM. To start the assay, 0.005 μM V_1_V_o_ and 0.3 mM ATP was incubated at room temperature (~ 25 °C) for 10 min with a volume of 100 μL. VopQ (1 μM) and the V-ATPase inhibitor folimycin (1 μM) were tested for ATPase activity inhibition. Reactions were stopped by adding 800 μL malachite green reagent followed by addition of 100 μL 34% (w/v) sodium citrate. The solutions were measured for absorbance at 620 nm in technical triplicates to detect free phosphate released from ATP hydrolysis. Assays were performed in at least three independent experiments. Clean pipette tips and microcentrifuge tubes were used in the sensitive assay to avoid background variation.

### Cryo-EM Image Collection

Cryo-EM grids were prepared using Vitrobot Mark IV (Thermo Fisher Scientific). Quantifoil R1.2/1.3 300-mesh gold holey carbon grids were coated with carbon film with thickness of 1~3 nm (Compact Coating Unit CCU-010, Safematic). 3.5 μL of protein sample was placed on a glow-discharged grid. After a waiting time of 15 s, the grid was blotted for 1.5 s under 100% humidity at 8 °C before plunged into liquid ethane. For grid screening, micrographs were acquired on a Talos Arctica microscope (Thermo Fisher Scientific) operated at 200 kV with a K3 direct electron detector (Gatan). The micrographs were recorded with SerialEM^34^. For data collection, micrographs were acquired on a Titan Krios microscope (Thermo Fisher Scientific) operated at 300 kV with a K3 direct electron detector, using a slit width of 20 eV on a GIF-Quantum energy filter. Images were recorded using SerialEM at a calibrated magnification of 59,983 with a super-resolution pixel size of 0.415 Å. The defocus range was set from −1.0 μm to −2.5 μm. Each micrograph was dose-fractionated (0.05 s/frame) with a total dose of around 60 e^−^/Å^2^.

### Cryo-EM Image Processing

Image processing was done roughly following the RELION-3.0 and RELION-3.1 user manuals^35^. The Titan Krios micrographs were motion corrected and binned two-fold with MotionCor2 (ref. ^36^), yielding a pixel size of 0.83 Å. The CTF parameters of the micrographs were estimated using CTFFIND 4 (ref. ^37^). All other steps of image processing were performed using RELION^35^. Around 1,000 particles were manually picked from a few micrographs for initial 2D classification and template generation for auto-picking. Particles were then autopicked from selected micrographs. After subgroup 2D and 3D classification with a binning factor of 2 or 4 for fast calculation, good class particles were selected for further processing. 3D classification with a binning factor of 1 allowed further separation of good particles from the bad and classification of different states. CTF-refinement was performed to improve the resolution of 3D reconstruction. Multi-body refinement was performed to analyze local regions with better map quality. The resolutions were estimated according to the gold-standard Fourier shell correlation (FSC) 0.143 criterion. RELION was used to calculate the local resolution map^35^.

Detailed workflows of image processing and statistics are shown in **Extended Data Figs. 1-3 and Extended Data Table 1**.

### Model Building and Structure Refinement

The structures of yeast V_1_V_o_ (PDB code: 3J9U)^7^ and V_o_-VopQ (PDB code: 6PE5)^17^ were used as initial models for model building of V_1_V_o_-VopQ (state2). The model was manually adjusted in COOT^38^. The structure of V_1_V_o_-VopQ was then used as initial model for V_1_V_o_ (state 1 and state 2). Structure refinement was performed using PHENIX^39^ in real space with secondary structure and geometry restrained. The statistics of the geometries of the models were generated using MolProbity^40^. The V_1_V_o_-VopQ model was validated without VopQ as the density for VopQ was not well resolved. Multibody refinement maps were merged in UCSF CHIMERA^41^ to generate a single map for validation of the overall models.

All structure figures were prepared with UCSF ChimeraX^42^ and PyMol^43^ and all EM density map figures were generated with UCSF CHIMERA^41^. Model building statistics are shown in **Extended Data Fig. 4 and Extended Data Table 1**.

## Data Availability

The atomic coordinates of the yeast V-ATPase complexes are deposited in the Protein Data Bank with accession codes 6XS0 (V_1_V_o_ state 1), 6XS1 (V_1_V_o_ state 2), and 6XS2 (V_1_V_o_-VopQ state 2). The corresponding electron microscopy density maps are deposited in the Electron Microscopy Data Bank with accession codes EMD-22298 (V_1_V_o_ state 1), EMD-22299 (V_1_V_o_ state 2), and EMD-22300 (V_1_V_o_-VopQ state 2).

## Supplementary Figures and Legends

**Extended Data Fig. 1.**
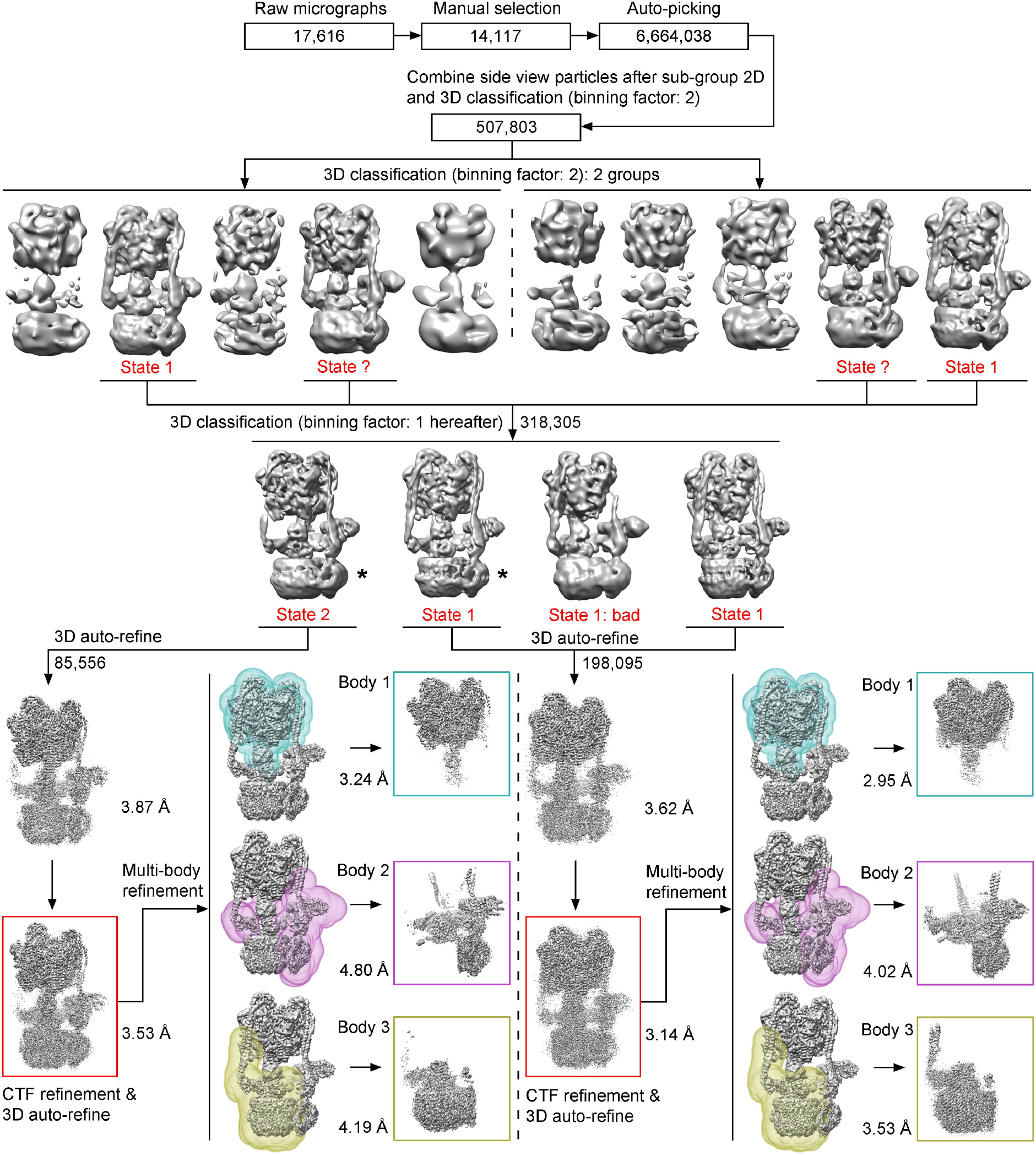
Flowchart of cryo-EM data processing for the yeast V_1_V_o_ holoenzyme. More details are described in methods and shown in **Extended Data Fig. 3**. 3D classes indicated with “*” are shown in **Extended Data Fig. 5a** in comparison with V_1_V_o_-VopQ cryo-EM density map.

**Extended Data Fig. 2.**
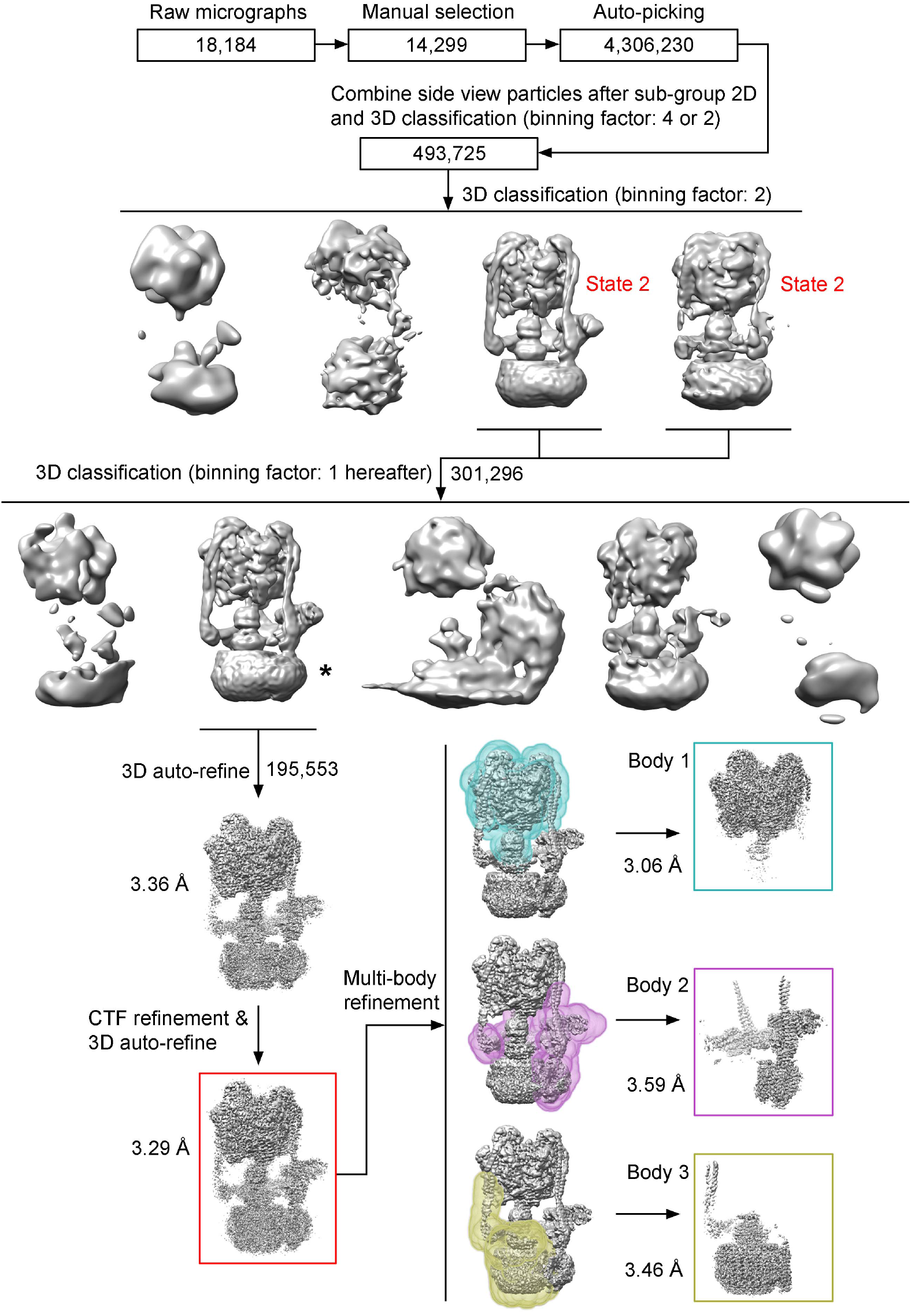
Flowchart of cryo-EM data processing for the yeast V_1_V_o_ holoenzyme in complex with VopQ. More details are described in methods and shown in **Extended Data Fig. 3**. The 3D class indicated with “*” is shown in **Extended Data Fig. 5a** in comparison with V_1_V_o_ cryo-EM density maps.

**Extended Data Fig. 3.**
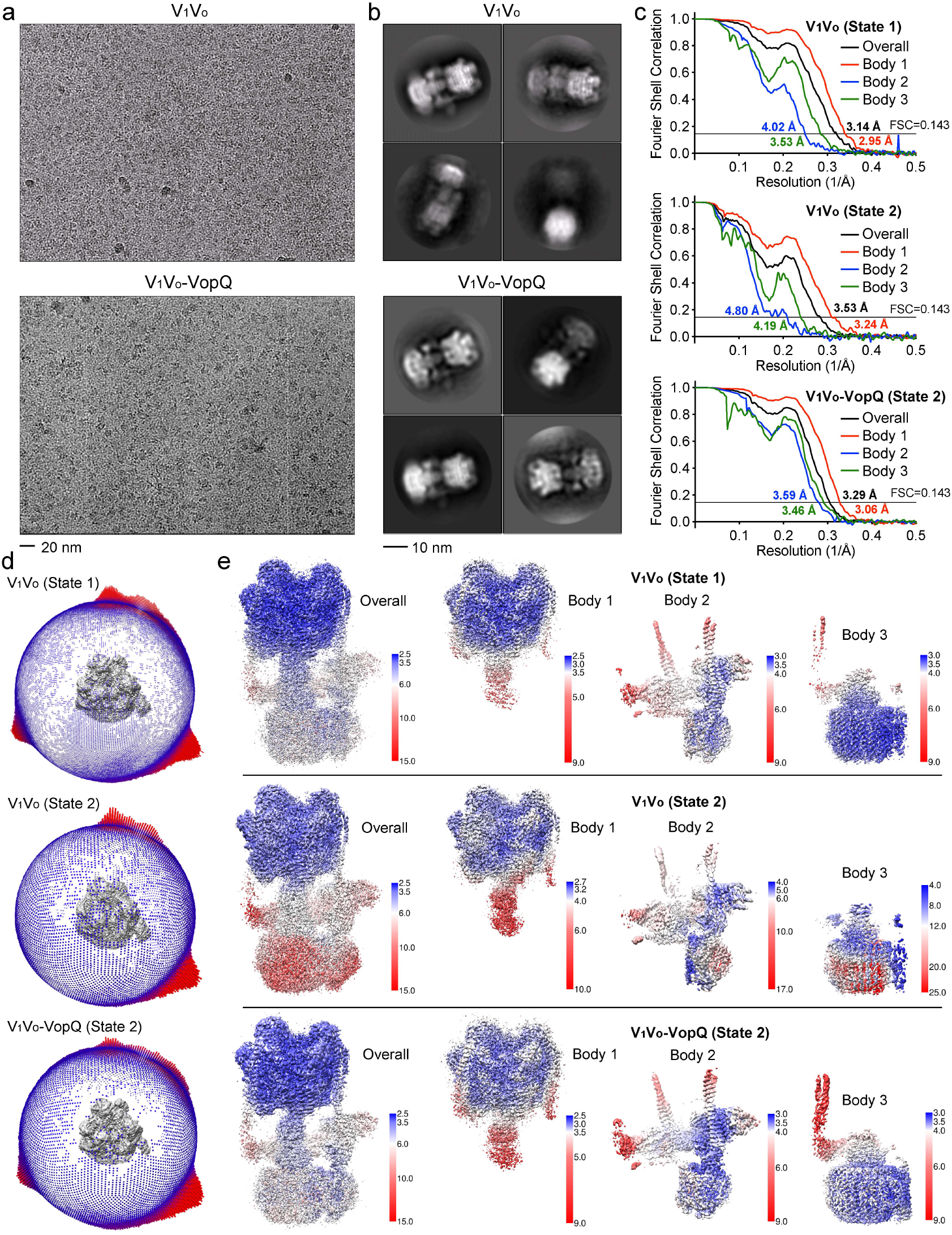
Analysis of the cryo-EM data. **a**, Representative cryo-EM micrographs. **b**, Representative 2D class averages. **c**, Gold-standard FSC curves for the cryo-EM density maps of the yeast V_1_V_o_ holoenzyme without and with VopQ bound. Multi-body refinement FSC curves are also presented. **d**, Angular distributions of particles used in the final 3D reconstructions, with the heights of the cylinders corresponding to the numbers of particles. **e**, EM density maps colored by local resolution.

**Extended Data Fig. 4.**
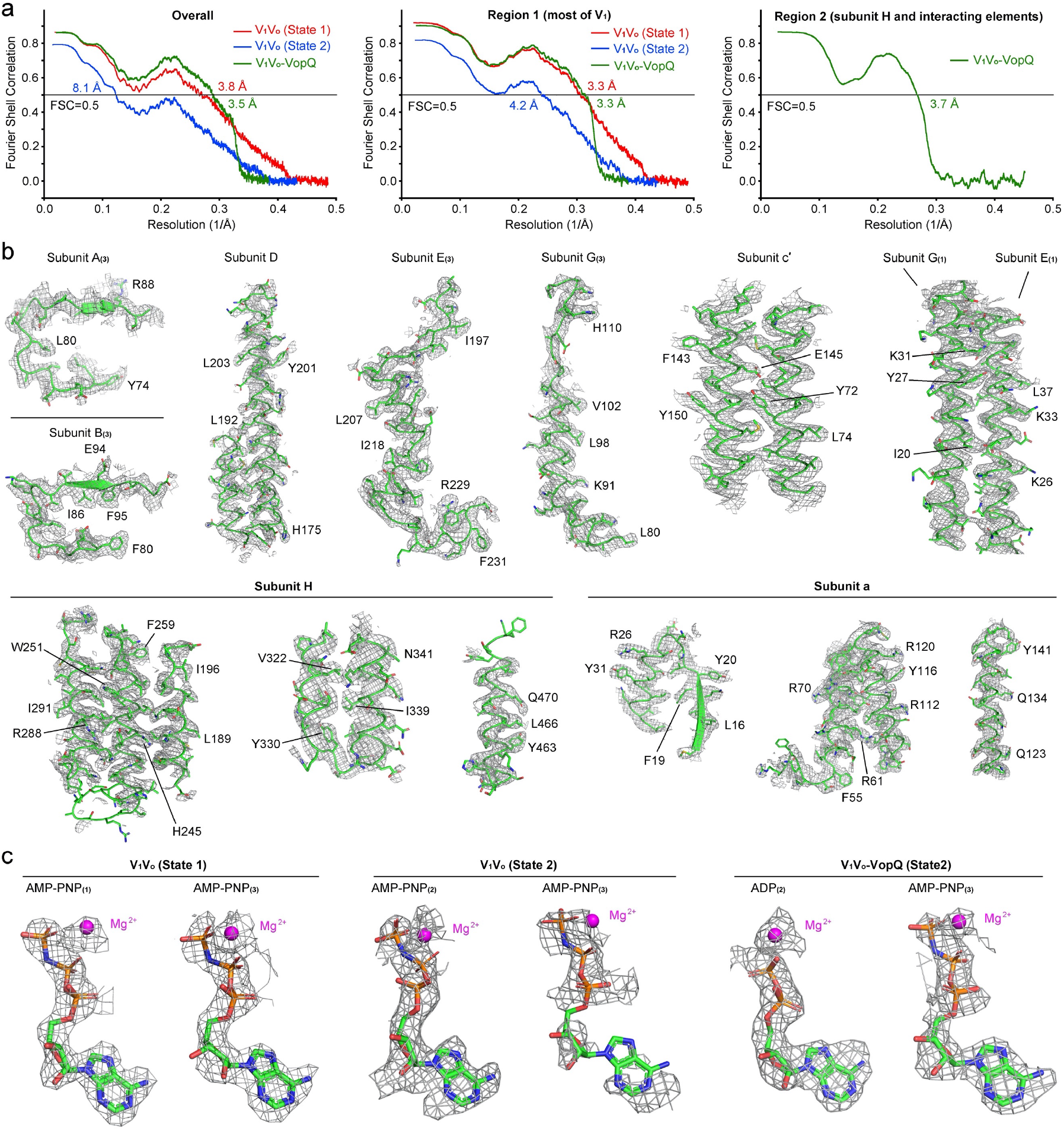
Model building of the yeast V_1_V_o_ holoenzyme without or with VopQ bound. **a**, FSC curves for cross-validation between the final refined models and the maps. Curves for the intact structure models (Overall) against the merged maps, for most of V_1_ subunits (Region 1 contains subunit A and B, 61-233 of subunit E, 61-114 of subunit G, 8-50_91-115_151-217 of subunit D, and ligands) against body 1 maps, for subunit H and neighboring subunits (Region 2 contains subunit H, 8-50 of subunit E, 2-40 of subunit G, and 3-380 of subunit a) against body 2 maps. The V_1_V_o_-VopQ model was validated (Overall) without VopQ as the density for VopQ was not well resolved. **b**, Local density maps shown as mesh for representative elements in the complex of V_1_V_o_-VopQ (contoured at 5.0 σ). Representative bulky residues are indicated. **c**, Local density maps for ligands of AMP-PNP and ADP (contoured at 10.0 σ).

**Extended Data Fig. 5.**
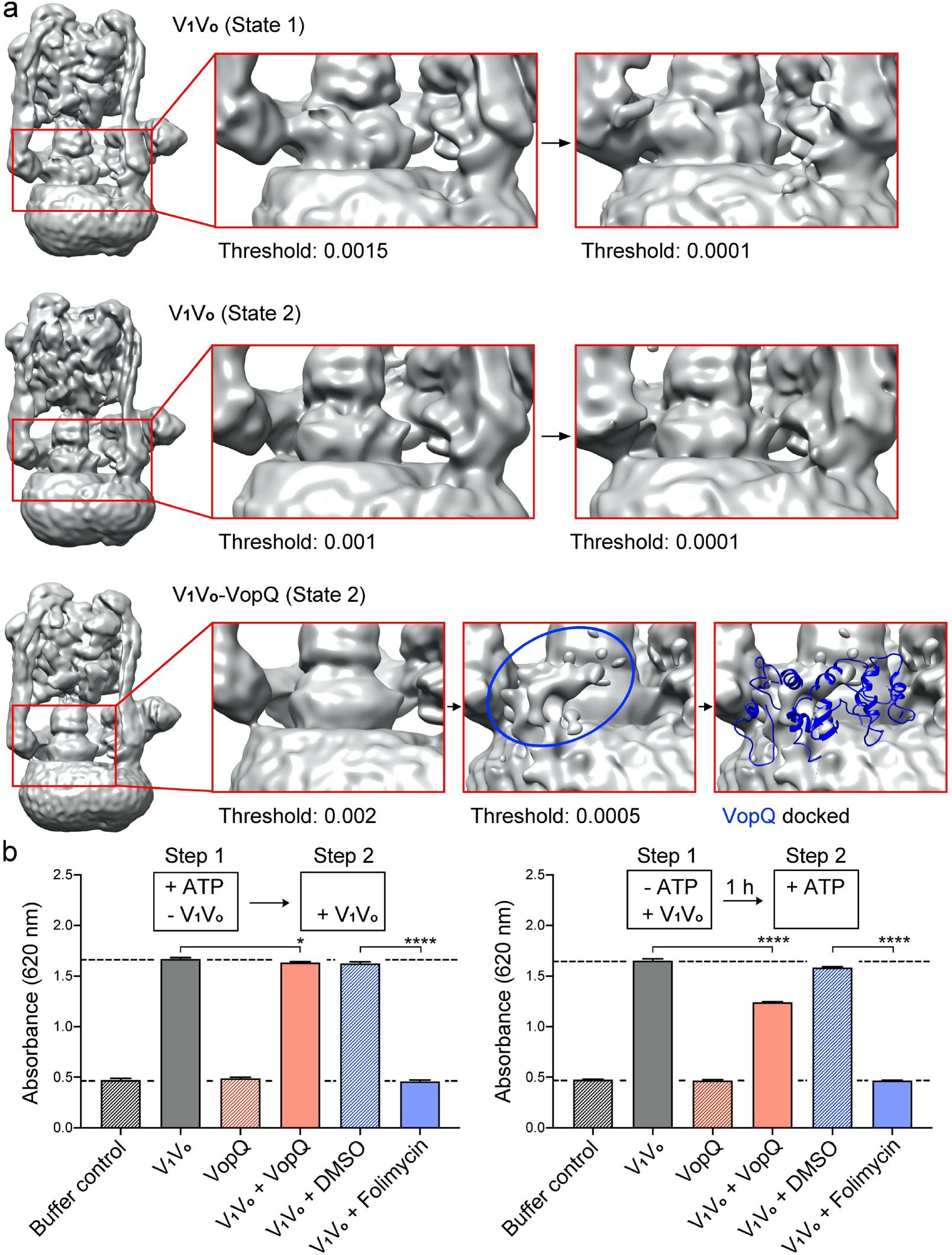
VopQ can bind the yeast V_1_V_o_ holoenzyme. **a**, Comparison of the 3D class cryo-EM density maps of the yeast V_1_V_o_ holoenzyme without and with VopQ bound. 3D classes indicated in **Extended Data Fig. 1-2** are shown at various thresholds. **b**, VopQ partially inhibits V_1_V_o_ ATPase activity *in vitro*. V_1_V_o_ was added into reaction systems with ATP present to initiate ATP hydrolysis (left), or without ATP to bind VopQ before ATP was added (right). Free phosphate produced from ATP hydrolysis was detected by malachite green assay. The V-ATPase inhibitor, folimycin, was used as a positive control. The baselines of buffer control and full activity are indicated. Data are displayed as mean with SD. Statistical significance was evaluated by using an unpaired Student t test (*P = 0.0106; ****P < 0.0001).

**Extended Data Fig. 6.**
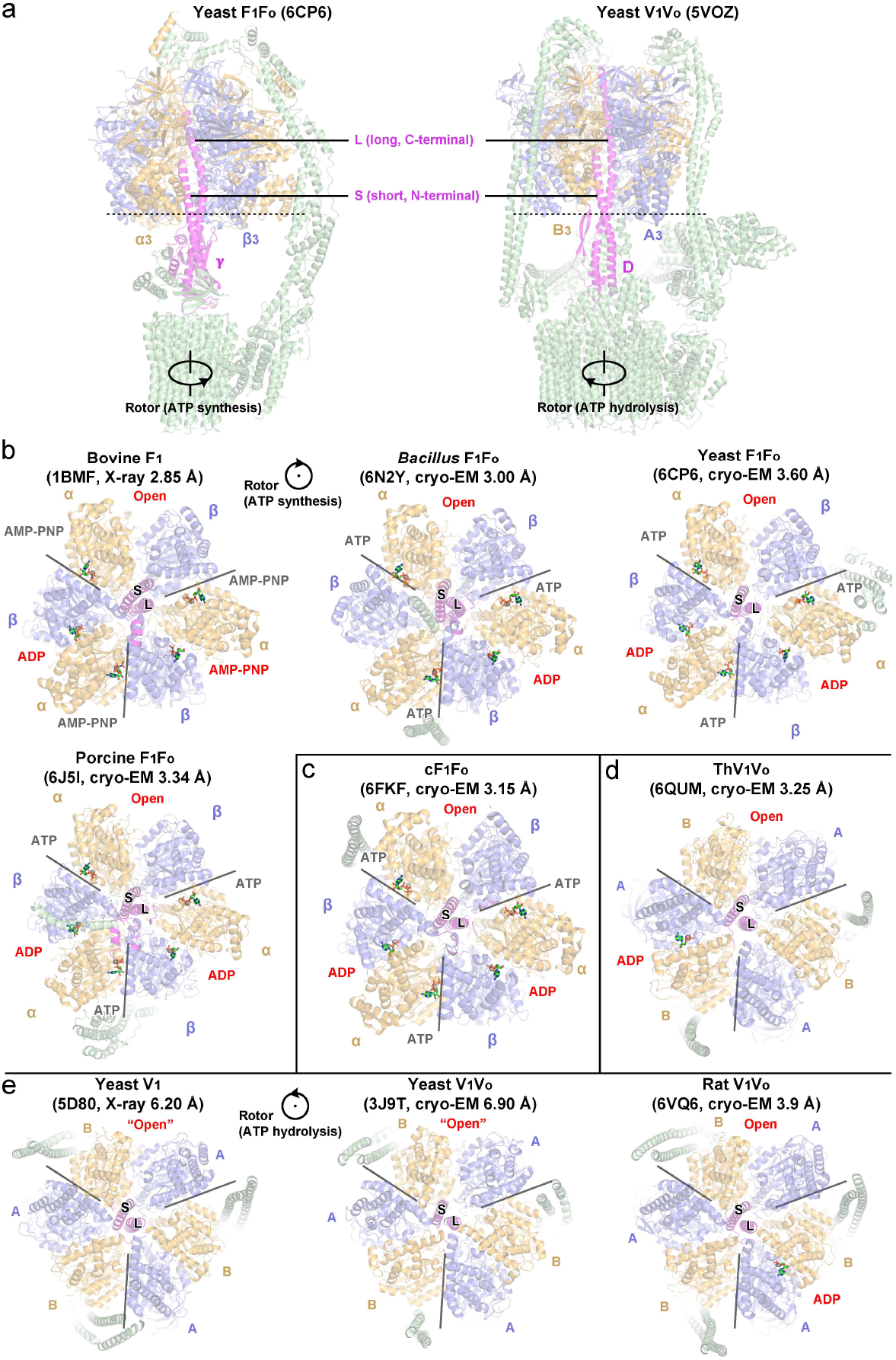
Representative conformations of rotary ATPases. **a**, Overall structures of a F-ATPase complex and a V-ATPase complex. Dash lines indicate cross section of catalytic subunits that are shown as bottom views in **b-e**. **b-e**, Bottom views of F_1_ and V_1_ catalytic subunits in representative conformations of F-ATPases (**b**), chloroplast F-ATPase (**c**), bacterial V/A-ATPase (**d**), and eukaryotic V-ATPases (**e**). The three catalytic subunit pairs (β-α in F-ATPases and A-B in V-ATPases) are indicated and separated by radial lines. Subunit γ or D and catalytic subunits are colored and indicated with all the other subunits in green. “S” or “L”, short N-terminal helix or long C-terminal helix of subunit γ and subunit D. Nucleotide molecules are shown as sticks with magnesium ions as spheres. Nucleotides at catalytic sites are labelled red, distinguished from those at non-catalytic sites (only in F-ATPases) in black. PDB accession code for each structure is indicated with the reported resolution noted. F-ATPase contains non-hydrolytic ATP molecules in F1 subcomplex between non-catalytic sites of α-β interfaces, but V-ATPase does not contain non-hydrolytic ATP molecules because of a major defect of P-loop in subunit B.

**Extended Data Fig. 7.**
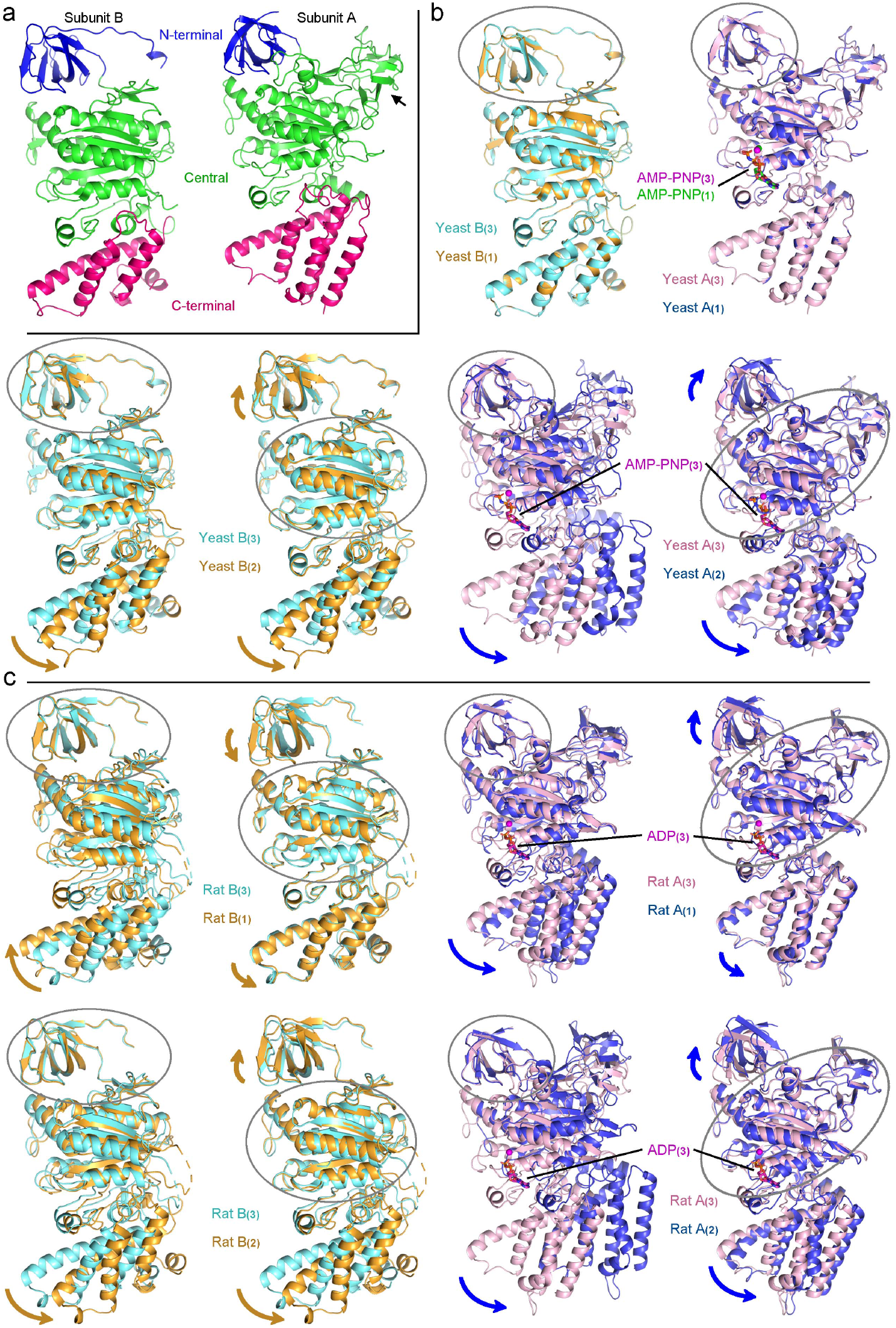
Comparison of the catalytic subunits in the yeast V_1_V_o_ complex (state 1) or the rat V_1_V_o_ complex (state1). **a**, Domain organization of the yeast catalytic subunits. Each domain is colored as indicated: N-terminal β-barrel domain (16-98 of B or 24-90 of A), central nucleotide-binding domain (99-390 of B or 91-475 of A), C-terminal domain (391-485 of B or 476-617 of A). Subunit A contains a small non-homologous domain (136-213) indicated by an arrow, which is for simplicity classified as part of the central nucleotide-binding domain. **b**,**c**, Comparison of subunit B and subunit A in the yeast and the rat V_1_V_o_ complexes. Subunit B_(3)_ or A_(3)_ is aligned against B_(1)_ or A_(1)_ and B_(2)_ or A_(2)_. The comparison is based on the N-terminal β-barrel domain or the central nucleotide-binding domain as indicated.

**Extended Data Fig. 8.**
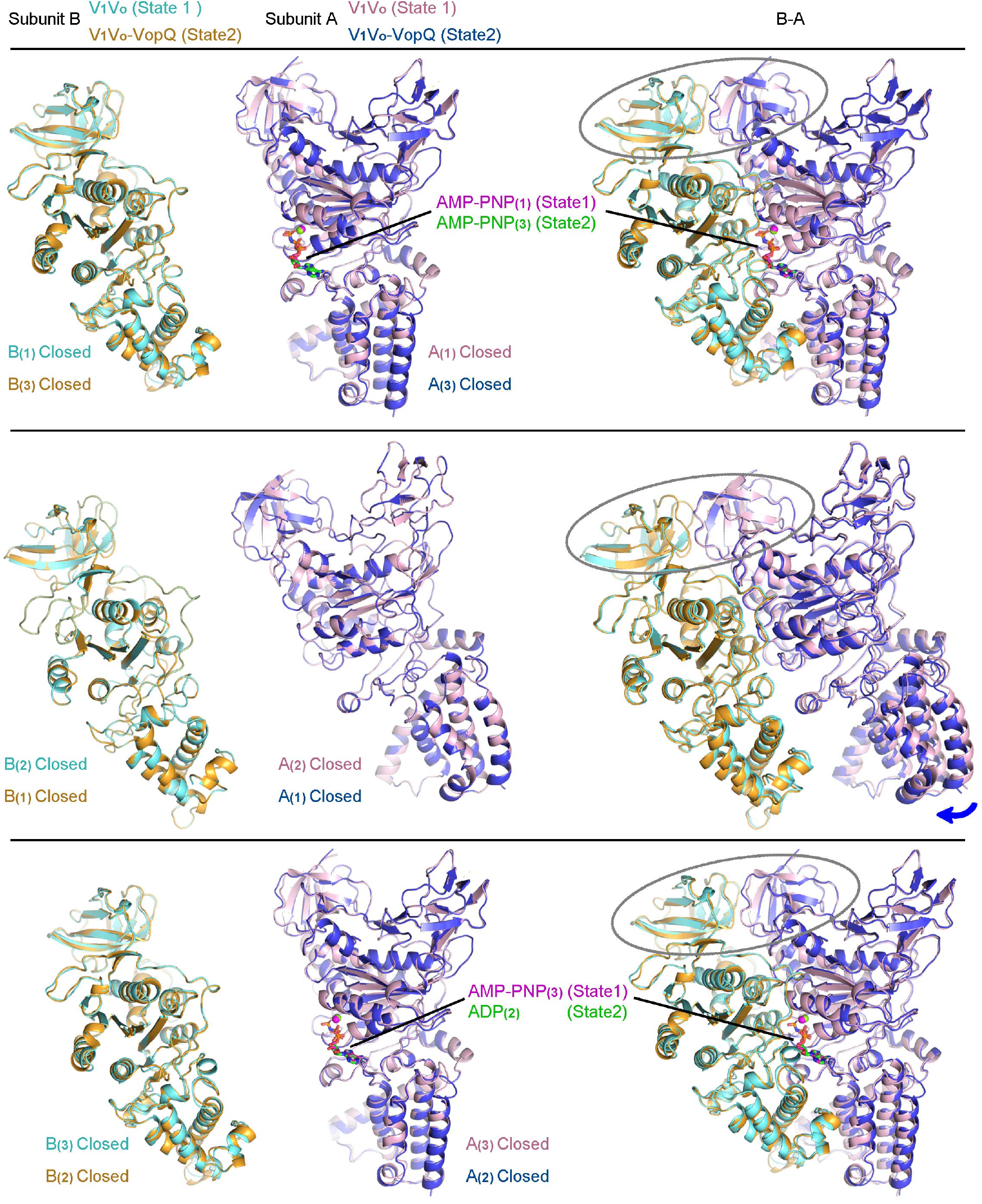
Comparison of the catalytic subunits in the yeast V_1_V_o_ complex (state 1) and the yeast V_1_V_o_-VopQ complex (state 2). Individual subunit B or subunit A is aligned against each other from the two complexes. A-B pairs are aligned against the N-terminal β-barrel domains as indicated.

**Extended Data Table 1.**
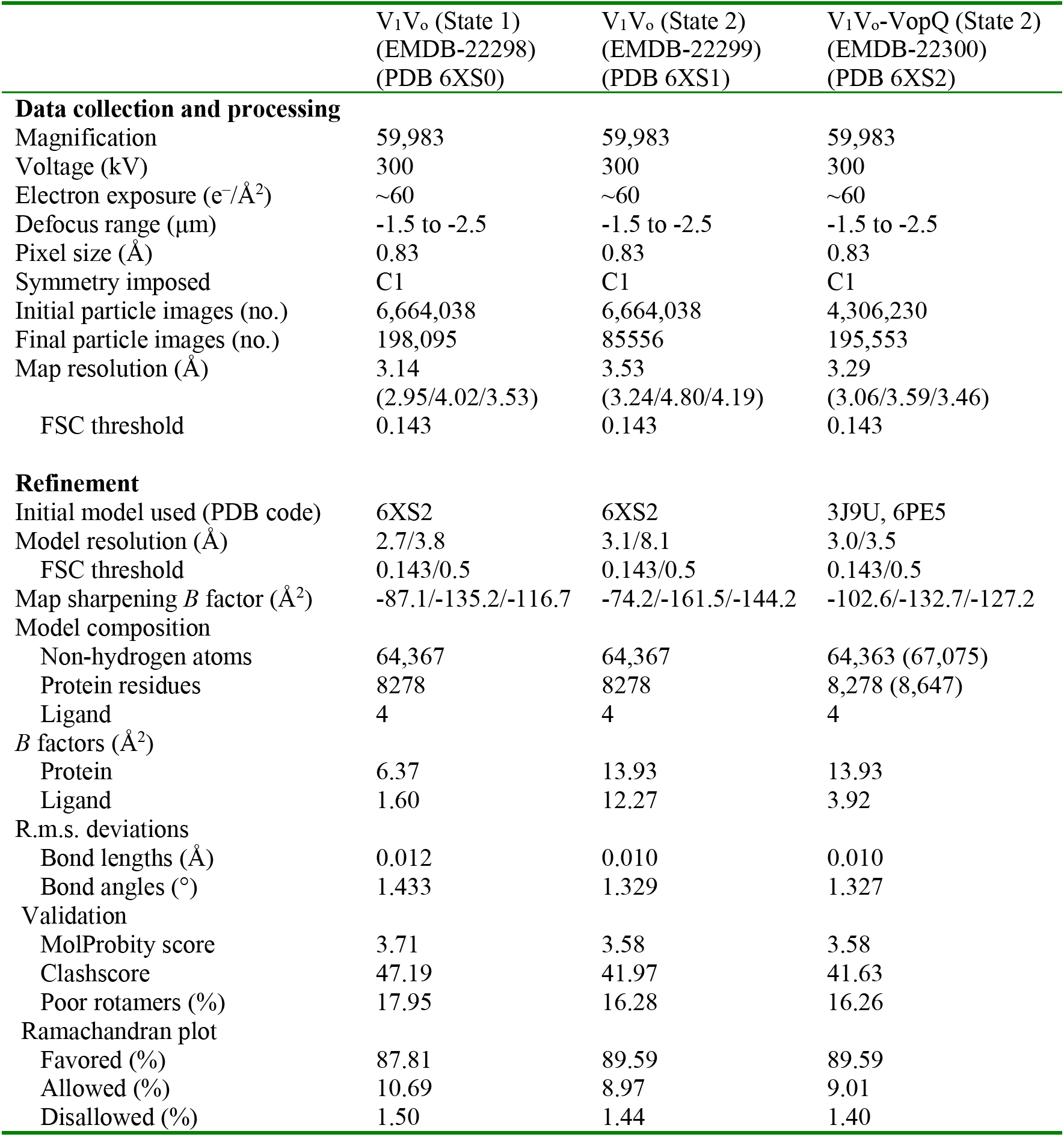
Cryo-EM data collection, refinement and validation statistics.

